# From Simulations to Inference: Using Machine Learning to Tune Patient-Specific Finite-Element Models of the Middle Ear Towards Objective Diagnosis

**DOI:** 10.1101/2024.10.15.618553

**Authors:** Hamid Motallebzadeh, Michael Deistler, Florian M. Schönleitner, Jakob H. Macke, Sunil Puria

**Affiliations:** Department of Communication Sciences and Disorders, California State University, Sacramento, CA, USA; Department of BioMedical Engineering, McGill University, QC, Canada; Machine Learning in Science, University of Tübingen, Germany; Tübingen AI Center, Tübingen, Germany; Neuroengineering – Elite Master Program, TUM School of Computation, Information and Technology, Technical University of Munich, Germany; Department Empirical Inference, Max Planck Institute for Intelligent Systems, Tübingen, Germany; Eaton-Peabody Laboratories, Massachusetts Eye and Ear; Speech and Hearing Bioscience and Technology, Harvard University Graduate School of Arts and Sciences

## Abstract

Computational models, particularly finite-element (FE) models, are essential for interpreting experimental data and predicting system behavior, especially when direct measurements are limited. A major challenge in tuning these models is the large number of parameters involved. Traditional methods, such as one-by-one sensitivity analyses, are time-consuming, subjective, and often return only a single set of parameter values, focusing on reproducing averaged data rather than capturing the full variability of experimental measurements.

In this study, we applied simulation-based inference (SBI) using neural posterior estimation (NPE) to tune an FE model of the human middle ear. The training dataset consisted of 10,000 FE simulations of stapes velocity, ear-canal (EC) input impedance, and absorbance, paired with seven FE parameter values randomly sampled within plausible ranges. The neural network learned the association between parameters and simulation outcomes, returning the probability distribution of parameter values that can reproduce experimental data.

Our approach successfully identified parameter sets that reproduced three experimental datasets simultaneously. By accounting for experimental noise and variability during training, the method provided a probability distribution of parameters, representing all valid combinations that could fit the data, rather than tuning to averaged values. The network demonstrated robustness to noise and exhibited an efficient learning curve due to the large training dataset.

SBI offers an objective alternative to laborious sensitivity analyses, providing probability distributions for each parameter and uncovering interactions between them. This method can be applied to any biological FE model, and we demonstrated its effectiveness using a middle-ear model. Importantly, it holds promise for objective differential diagnosis of conductive hearing loss by providing insight into the mechanical properties of the middle ear.

## Introduction

Computational models have been used to explain experimental data, as well as to interpret and predict system behavior under circumstances that, due to measurement limitations, may be difficult to study experimentally. One of the challenging aspects of developing valid models, particularly finite-element (FE) models, is the potentially large number of parameters that need to be specified. In this study, we implemented a machine-learning approach to tune FE model parameters against experimental measurements. This approach is generally applicable to any computational modeling method; however, the focus of this work was on a human middle-ear (ME) model, a popular model in biomechanics of hearing. In contrast to lumped-element models, FE models are built upon realistic geometrical and physical representation of the biological system being simulated, enabling explicit mapping of model parameters to ME anatomy and physiology (e.g., Gan et al., 2002; O’Conner et al, 2016).

To develop FE models, the geometry is typically reconstructed by segmenting CT, or for higher spatial resolution using micro-CT images. After geometry creation and meshing, physical parameters are assigned. Due to various factors, a wide range of parameter values have been reported. For example, the eardrum’s Young’s modulus values range from 2 MPa (Aernouts et al., 2012) to 300 MPa (Fay et al., 2005) depending on how the sub layers were treated. Even more challenging is that unknown parameters are often adopted from other specimens, or similar tissues (Motallebzadeh and Puria, 2021). Other complexities in defining material parameters include anisotropy (Fay et al., 2006), viscoelasticity (Motallebzadeh et al., 2015), and material and geometric nonlinearities (Motallebzadeh et al., 2013). Thus, material model selection depends on multiple factors including simulation objectives, computational cost, and parameter availability.

Assigning boundary conditions and extracting appropriate data from models are additional steps that can significantly impact computational study outcomes. For instance, simplifying the eardrum extension by clamping its annular ligament affects TM behavior (O’Connor et al., 2017). Similarly, the ME cavity, with its complex air-cell structure, is often modeled as a simple smooth, loss-less hard wall (Keefe, 2015; Motallebzadeh et al., 2017a). Data extraction methods in FE models must mimic experimental conditions. For example, experimental vibrometry often uses a microphone tube with a 2-mm from the umbo (Khaleghi et al., 2015), whereas FE models typically average pressure over the entire ear canal (EC) surface (Nørgaard et al., 2024). Measurement trajectory uncertainties, especially in vibrometry, arise due to experimental limitations leading to non-orthogonal measurement directions that can extenuate certain modes of vibrations (Dobrev et al., n.d.; Khaleghi et al., 2016).

In conventional procedures, model parameters are tuned through systematic one-by-one sensitivity analyses, iteratively comparing simulation data with experimental measurements (e.g., De Greef et al., 2017). Such an approach to sensitivity analyses often lacks comprehensiveness, failing to explore potential parameter interactions between multiple parameters (Maftoon et al., 2015). In addition, empirical data is often represented as a population average, where fine features of individual responses can be masked.

As a result, tuning computational models has generally been subjective, aiming for a general agreement with averaged empirical data characteristics, such as resonance frequency and high-frequency corner frequency (Motallebzadeh et al., 2017a). This process becomes even more complex when models attempt to replicate multiple independent characteristics, like eardrum vibrational modes and continuous impedance spectrum. In addition, developing and tuning a reliable model can take several years. For instance, one of our comprehensive ME models (O’Connor et al., 2017) was refined over a period of 6-7 years, starting from earlier work (Cai et al., 2012) and further developed through intermediate iterations (O’Connor et al., 2016).

There have been examples of semi-automatic optimization procedures to tune the mechanical parameters of individual ME parameters (Aernouts and Dirckx, 2012; Motallebzadeh et al., 2013). Even those methods were limited to tuning only two parameters within much simpler models consisting of a single component (just the eardrum). Sackmann et al. (2019) introduced inverse fuzzy arithmetic for parameter identification in model-based approaches to detect pathological conditions of the ME. Their technique, a possibilistic method, which assesses the plausibility of outcomes based on imprecise or incomplete data, is suitable for stochastic uncertainty analysis. In another study (Sackmann et al., 2022), they selected multiple fitting criteria, such as resonance characteristics, and used averaged sensitivity indices to identify parameter values for a simplified ME model based on reference experimental data (Merchant et al., 2016). These methods have been applied to classify ME pathologies (Winkler et al., 2024). However, normal intersubject variabilities in ME structures (Bartling et al., 2021) can be misinterpreted by machine-learning algorithms trained on discrete simulators.

Simulation-based machine-learning methods are emerging as a powerful tool for interpreting clinical tests. Here, we propose using simulation-based inference (SBI) (Cranmer et al., 2020; Tejero-Cantero et al., 2020) to identify parameters in biological models that can reproduce reference experimental data. Specifically, we employ the Neural Posterior Estimation (NPE) method (Papamakarios and Murray, 2016). In NPE, parameter sets are initially drawn from a prior distribution, with parameters sampled randomly within the specified plausible range (Greenberg et al., 2019). These parameters are then run through a model (in this study, a FE model) to generate pairs of input parameter values and corresponding simulation outputs. NPE trains a deep neural density estimator to predict parameters from simulated data. This process allows one to effectively incorporate the knowledge gained from having already developed a valid FE model into the NN with known input output relationships. This is more efficient than making massive amounts of physiological or clinical measurements where the input-output relationships are not well characterized. The detailed procedure of SBI and the architecture of its neural network can be found in our previous publications (Greenberg et al., 2019; Papamakarios and Murray, 2016; Tejero-Cantero et al., 2020).

NPE offers two key advantages over previous methods: first, it provides a principled way to handle noisy measurements. Unlike methods such as genetic algorithms, NPE infers the Bayesian posterior distribution given experimental data, capturing any parameter settings consistent with noisy data. Second, the method is amortized; once trained, it can rapidly infer parameters from new measurements without requiring new simulations or retraining. This enables the determination of ME parameters for both normal and pathological ME which has applications in clinical settings.

## Methods

First, we introduce the pipeline for the SBI methodology. Then, the experimental procedure for obtaining the reference empirical data used to tune the model is detailed, along with a summary of the FE model development procedure and prior knowledge of the model parameters. The final section explains the procedure for running a large number of simulations, extracting data, and preparing it to train the SBI.

### A. SBI Methodology

Figure 1 illustrates the typical flow of the SBI method. After developing the simulator and selecting parameters of interest, a plausible range for each parameter is chosen based on literature data (Fig. 1A). For missing parameter values, plausible ranges are assigned from morphologically similar tissues (Motallebzadeh et al., 2017b, 2018). Parameters are randomly sampled within these ranges and fed into the mechanistic model (Fig. 1B). Simulation data are collected (Fig. 1C), and optionally, a pre-processing step reduces the dimensionality of the simulation results. Each input parameter set and the corresponding simulation outcome are then paired and fed into a neural network. The neural network learns to associate input parameters with the simulation outcomes (Fig. 1D). Once trained, the neural network can predict the range of parameters consistent with the reference data (Fig. 1E) in the form of a probability distribution over parameters (the posterior distribution, Fig. 1F) when the reference measurement data is used as an input.

**Figure 1.**
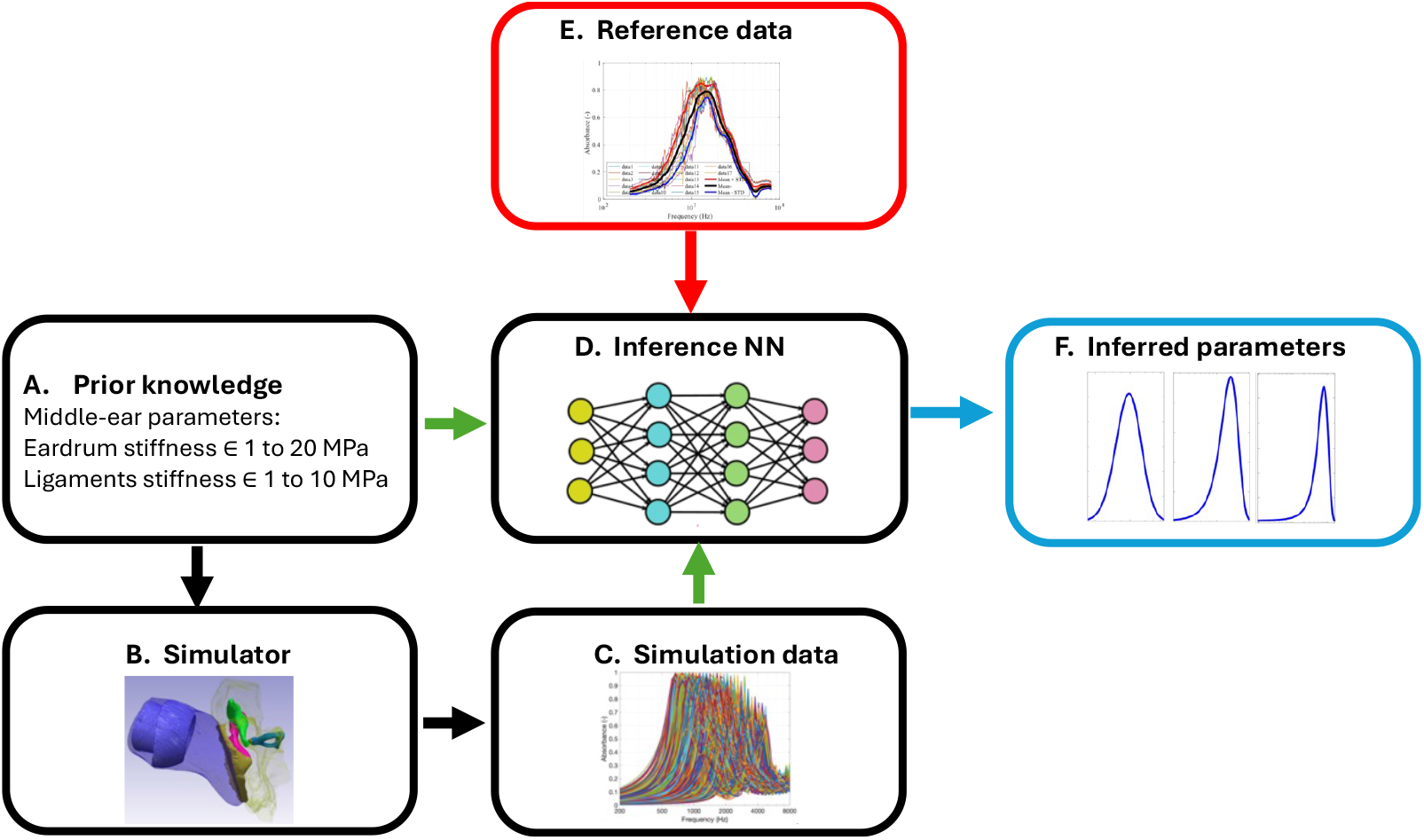
Schematic flow of SBI: (A) Prior range of simulator parameters are identified, and a subset of random values within this range selected and (B) imported into a finite element model termed mechanistic ‘simulator’ to generate the (C) simulation data. Multiple simulations are performed. (D) The paired sets of simulator parameter inputs and simulation data outputs are then introduced to the SBI neural network (NN), which learns their input-output relationships and that effectively trains a NN. (E, F) When a reference data set (measurements) is introduced, the NN infers the probability distributions of the parameters that would have produced that data.

### B. Imaging and physiology measurements

The reference experimental measurements for this study were obtained from two independent sets of experiments. First, is wideband tympanometry which provides a measure of the input impedance of the ME. This allows for the calculation of ME absorbance, indicating the ratio of absorbed energy by the ME to the input acoustic energy. The second is the stapes velocity, which describes the transmission through the ME. Details of the measurement procedure can be found elsewhere (e.g., Sim et al., 2010). Seventeen sets of stapes velocity, wideband impedance, and absorbance spectra at ambient pressure were collected from test-retests on a single temporal bone. These measurement data were sourced from another project in our lab (Dharmarajan et al., 2020).

After conducting the measurements, the temporal bone specimen was taken to Boston University, where μCT imaging was performed (Zeiss, Xradia 520 Versa) at a resolution of 15.1 um. The image stack was imported into Simpleware (Version 2019; Synopsys, Inc., Sunnyvale, USA), and a combination of manual and automatic segmentations was used to reconstruct the 3D geometry of the ME of that specimen (Fig. 1B). The model includes the EC, eardrum, ossicles, ossicular joints, and suspensory ligaments and tendons. For the experimental condition, the ME cavity was open and thus excluded from the model to reduce computational cost. The model input load was applied as a prescribed volume velocity at the probe-EC interface. Simulations were performed at 50 frequencies, logarithmically spaced between 250 and 8,000 Hz.

To calculate impedance, the ratio of averaged pressure and volume velocity of the EC air was extracted on a circular plane with a radius of 3 mm at the center of the EC, 3 mm from the probe-EC interface, comparable to the distance between speaker and microphone in the tympanometer probe used in the experiments (Keefe et al., 1993). The EC input impedance (*Z*_*ec*_) is the ratio of the pressure over the volume velocity (Motallebzadeh and Puria, 2021). The absorbance, the ratio of the acoustic stimuli energy absorbed by the ME can be calculated from the impedance by

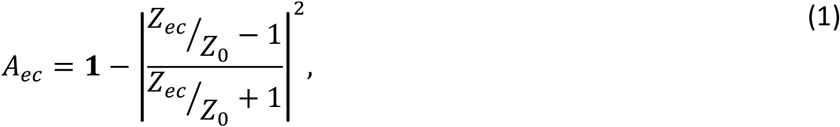

where, *Z*_0_ = ^*ρc*^/_*a*_ corresponds to the input impedance of an infinitely long tube. In this equation, *ρ* and *c* are the respective density of air and speed of sound in air, and *a* is the lateral EC area.

The normalized stapes velocity was calculated as the ratio of the stapes velocity in the piston-motion direction to the pressure (V_st_/P_ec_), measured approximately 2 mm from the umbo.

For the two experimental measurements— tympanometry and stapes velocity —the exact location of the measurement probe was not readily accessible, introducing a degree of uncertainty. To address this, we conducted a series of analyses to pinpoint the reference measurement location for the FE model. This was achieved through bulk simulations of the training dataset, as detailed in Supplementary Information (SI-1). We found that averaging pressure on the plane near the eardrum, parallel to the probe surface, yielded stapes velocity, EC impedance, and absorbance values consistent with experimental data. Without this approach, parameter variation alone failed to match the experimental data, preventing SBI from converging to provide a probability distribution of the parameters.

### C. Prior parameter knowledge and the finite element model

We selected seven ME parameters to tune our model: Young’s moduli of the eardrum, mallear ligament, stapes annular ligament (SAL), ossicular joints, damping coefficient, and cochlear load. These parameters were chosen because they more significantly influence ME behavior and because they are often associated with ME pathologies. For instance, stiffening of the mallear ligament is linked to ossicular fixation while SAL is linked to otosclerosis. Plausible ranges of these parameters were selected from experimental data reported in the literature for normal MEs (Table 1). Parameters with less uncertainty and/or sensitivity, such as the Young’s modulus of ossicles (∼14 GPa), density of soft and hard tissues (1100 and 2100 kg/m^3^, respectively), and Poisson’s ratio of soft and hard tissues (0.485 and 0.3, respectively), were assigned from the literature (Motallebzadeh et al., 2015, 2017b).

**Table 1.**

| Table 1 |  |  |  |
| --- | --- | --- | --- |
| Material properties |  |  |  |
|  | Minimum | Maximum | Reference(s) |
| Young’s modulus (MPa) |  |  |  |
| Eardrum | 1 | 20 | (Aernouts et al., 2012; Decraemer et al., 1980) |
| Stapes annular ligament | 0.005 | 1 | (Gan et al., 2011; Kwacz et al., 2015) |
| Mallear ligament | 1 | 10 | (Cheng and Gan, 2008a) |
| Soft tissue (suspensory ligaments and tendons) | 1 | 10 | (Cheng and Gan, 2008a, 2008b) |
| Joints | 1 | 10 | (Zhang and Gan, 2011) |
| Damping (-) | 0.05 | 0.2 | (Zhang and Gan, 2010) |
| Cochlear load (GΩ) | 0.15 | 1.5 | (Frear et al., 2018) |

The FE simulations were performed in COMSOL Multiphysics® software version 6.2 following the ‘Method’ procedure (CMSOL Blog, 2017), with data were directly exported following the procedure explain in (COMSOL Blog, 2017) to accelerate the large simulations. We automated the randomized selection of parameter values within the prior ranges, imported each set into COMSOL, conducted parallel simulations, and processed the data using a MATLAB script (Version: 9.14.0 (R2023a)). We performed 10,000 simulations simultaneously on two computers using a hybrid shared and distributed memory method (Frei, 2014). The simulations took about four weeks, with each run taking 260 seconds for 50 frequencies.

### D. Training the neural network

We ran SBI with the 10,000 parameter values paired with corresponding FE model simulation outputs, which included stapes velocity and ME impedance (both magnitude and phase) as well as absorbance magnitude, concatenated into a single vector of 250 elements. Gaussian noise was then added to this concatenated vector, with the standard deviation for each sub-vector—stapes velocity, ME impedance, and absorbance—applied in the order of their inclusion, based on the variability extracted from experimental measurements. This ensures that the NN is exposed to and learns from the diversity of the experimental data. The parameters and simulated traces were z-scored before being fed to the neural network. The neural network used is a conditional density estimator, specifically a conditional neural spline flow (NSF) employing a fully-connected embedding network with three layers and fifty hidden units each (Durkan et al., 2020).

## Results

Initially, we evaluated the performance of the trained SBI on FE simulation data with predefined random parameters (Fig. 2). Subsequently, we employed the trained NN to infer the parameters of the baseline FE model using reference experimental data (Fig. 3). We conducted a sensitivity analysis on the additive noise level to evaluate the robustness of the SBI performance (Fig. 4). Additionally, we investigated the effect of training dataset size (Fig. 5).

**Figure 2.**
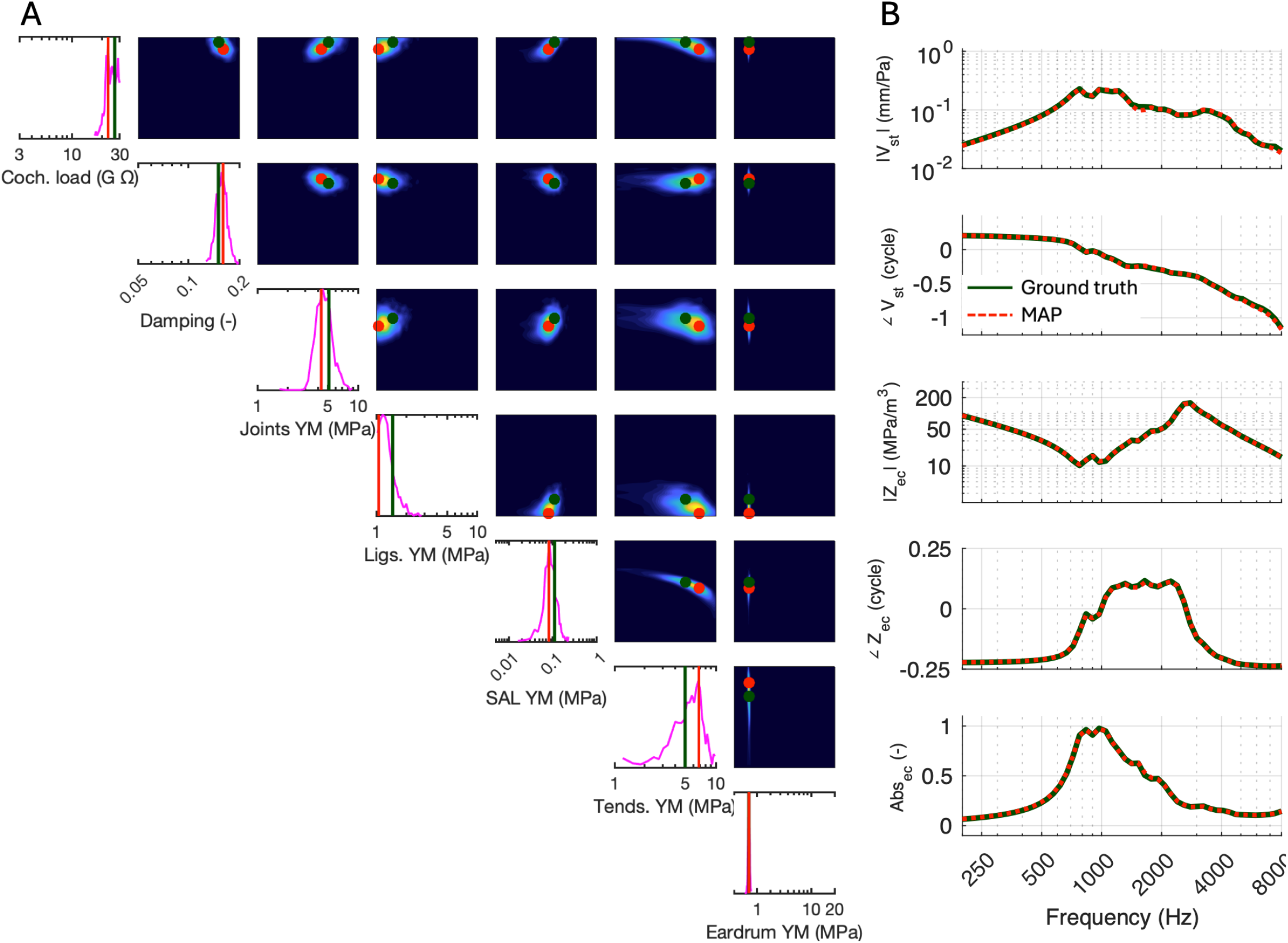
SBI Validation: The SBI baseline neural network (NN) was trained with 10,000 simulations. A new set of random parameter values (represented by vertical dark green lines and dots in A) was input into the FE simulator. The simulation results for the magnitude and phase of the stapes velocity (Vst/Pec), impedance (Z_ec_), and the magnitude of the absorbance (A_ec_) calculated at 50 frequencies (depicted as dark green lines in Fig. B) were fed into the trained NN. The NN returned probability distributions for each parameter in 1D (shown as magenta histogram lines in the diagonal panels of A) and paired histograms, represented as confusion matrices (off-diagonal color maps in A). The NN also returned the maximum *a posteriori* (MAP) values for each parameter (indicated by red vertical lines and dots in A). These MAP values were then imported into the FE simulator, and the resulting stapes velocity, impedance, and absorbance were plotted and are shown as red dotted lines, in B and overlap with the original data (black lines) used to estimate the parameter distributions.

**Figure 3.**
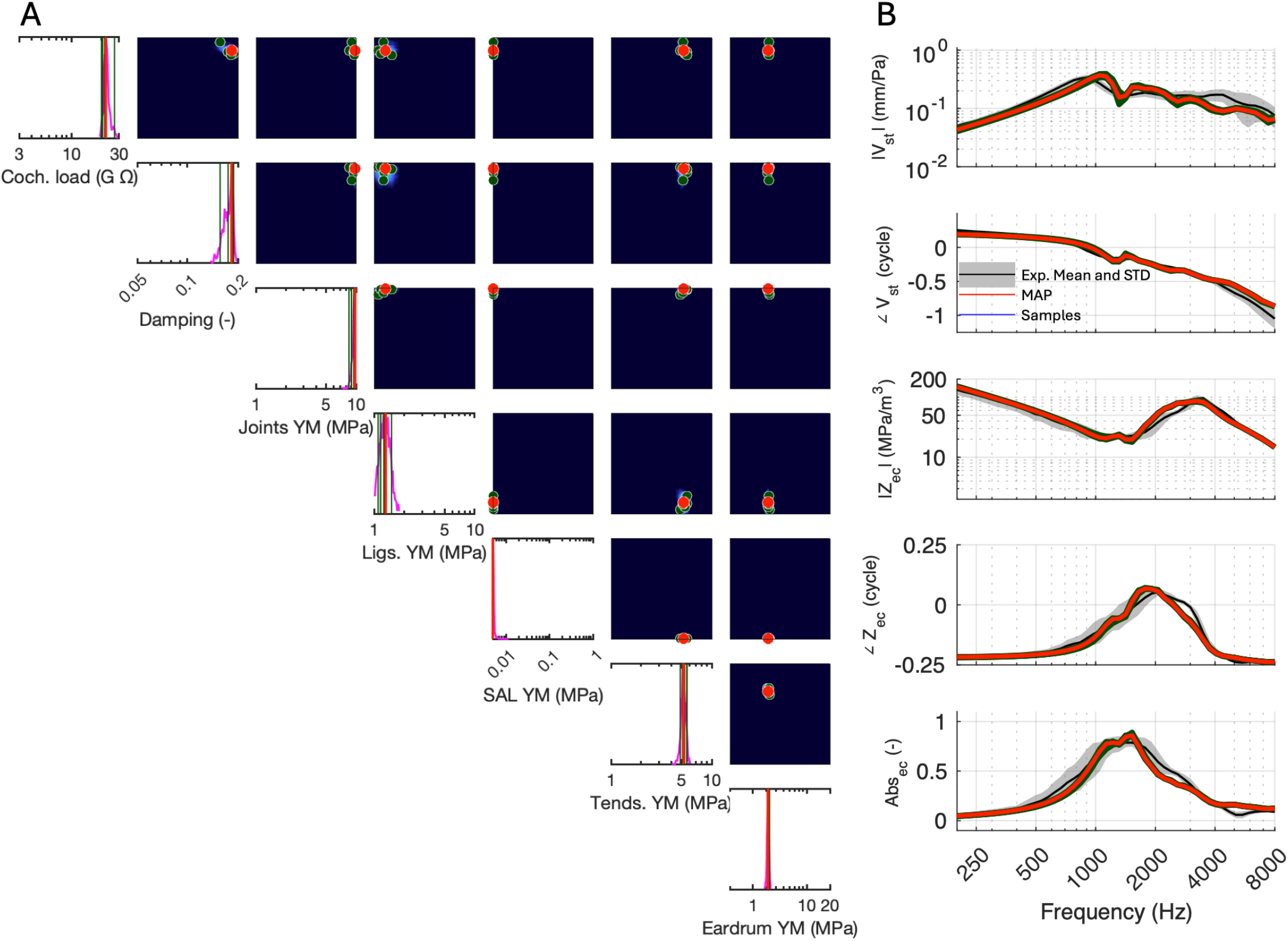
Inferring the model parameters from reference experimental data. Reference data averaged over 17 test-retest data from a single temporal bone was used to determine parameters from the initially trained NN of Figure 2. (A) For the given input data set, the NN returned the MAP (vertical red lines and dots) and five arbitrarily chosen parameter sets (vertical dark-green lines and dots) within the probability distribution of the parameters (magenta curves), which are then imported into the FE simulator and results calculated. (B) The resultant spectra (red line) of the MAP and for the 5 samples (dark-green lines) are compared with the objective experimental data (black curves).

**Figure 4.**
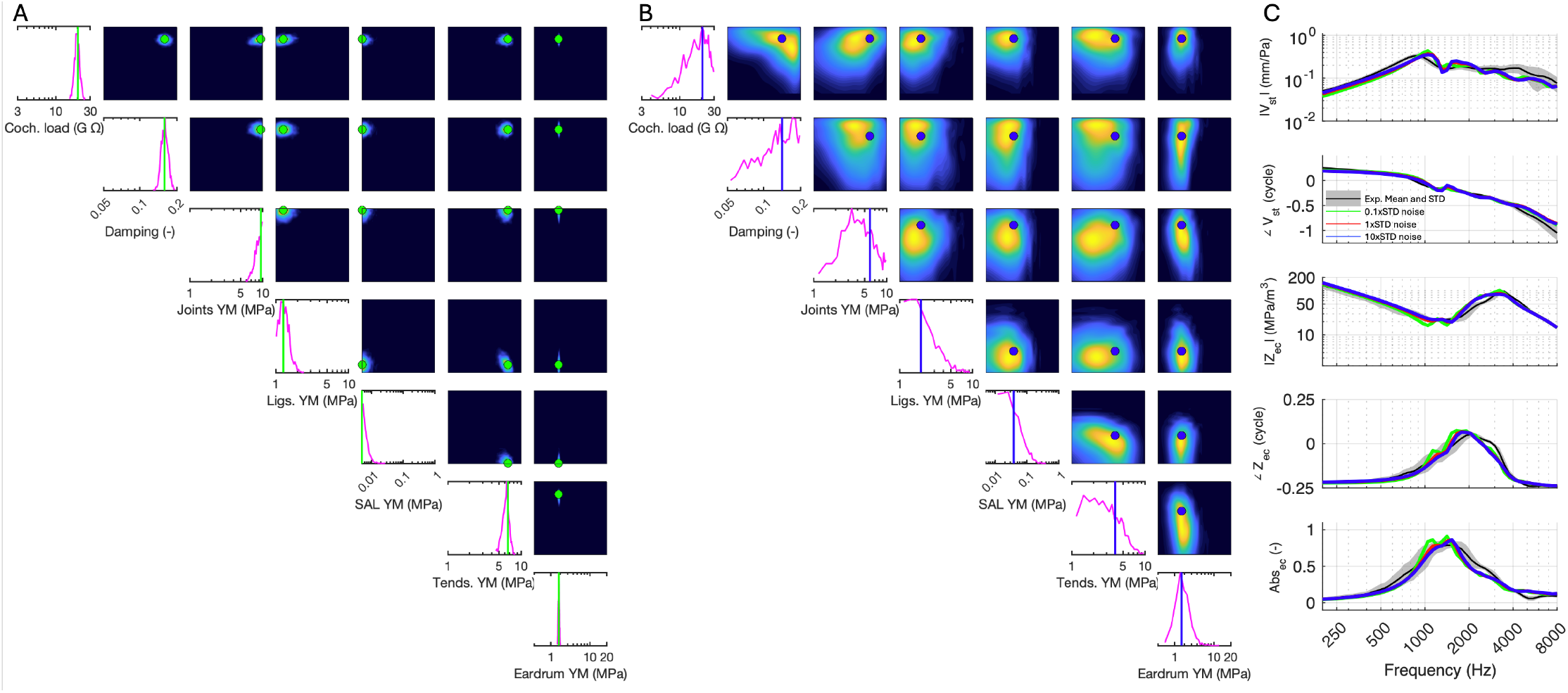
The effect of added noise to the training data on NN performance: The baseline noise (1x STD) was extracted from the experimental data and added to the simulation data obtained from the FE simulator, then magnified by factors of 0.1, 1, and 10. These modified data sets were imported into the NN for training. The three trained NNs were then each exposed to the experimental data, returning the probability distributions of the parameters for (A) 0.1xSTD, 1xSTD, and (B) 10xSTD. The descriptions of the figures and the NN output for 1xSTD noise are presented in Figure 3. (C) The MAP values of the predicted distributions were imported into the FE model, and the results are compared with the experimental data.

**Figure 5.**
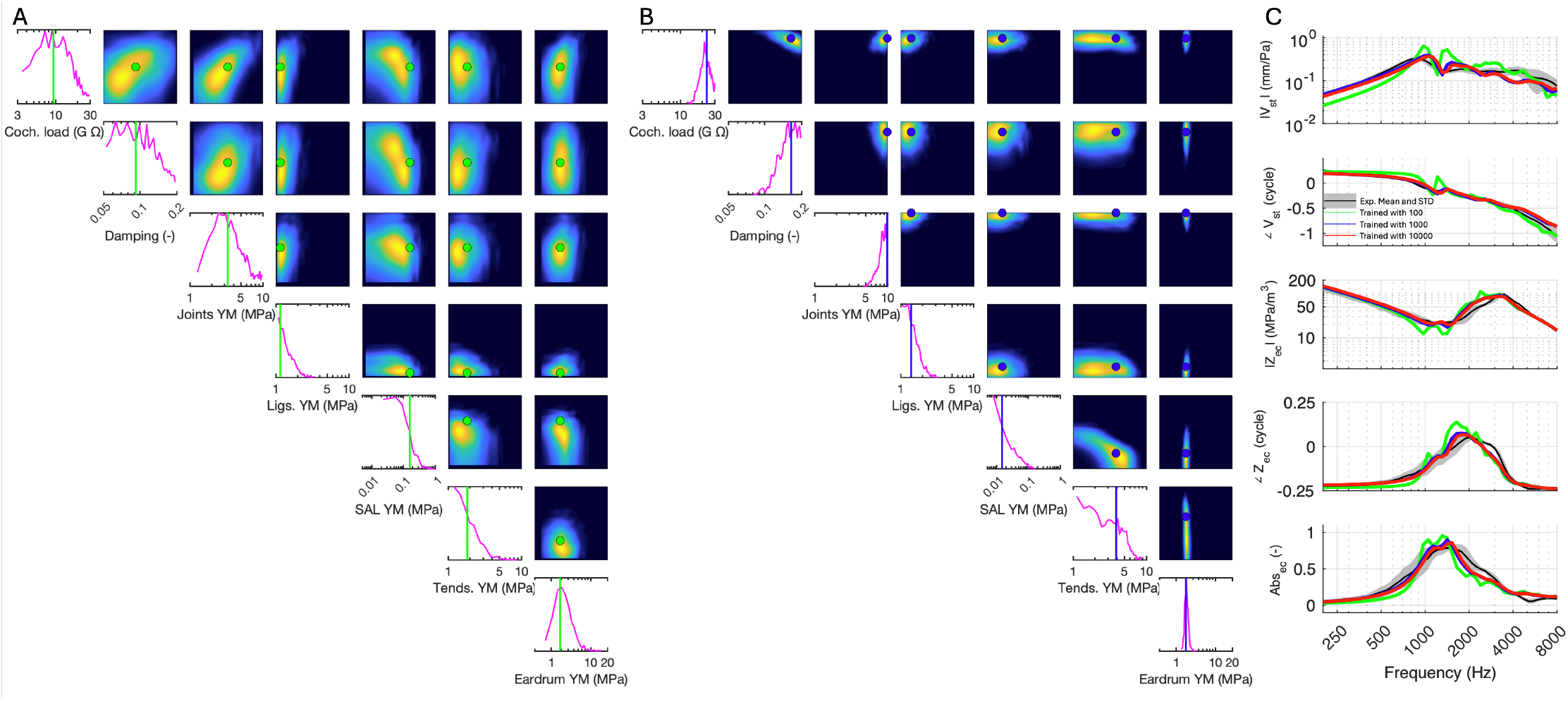
Effect of training dataset size on NN performance: The NN was trained with 100, 1,000, and 10,000 datasets. The three trained NNs were then exposed to the experimental data, returning the probability distributions of the parameters for (A) 100 and (B) 1,000 FE model simulations. The results for 10,000 model simulations and the panel descriptions are presented in Figure 3. (C) The MAP values of the predicted distributions were imported into the simulator, and the resultant spectra were compared with the experimental data.

### A. Neural Network training and validation

For preliminary work we performed 10,000 simulations, each containing a set of prior parameter values paired with corresponding simulation results. The simulation results include the magnitude and phase of the stapes velocity (V_st_/P_ec_), impedance Z_ec_, and the magnitude of the absorbance (Abs_ec_) calculated at 50 frequencies logarithmically spaced from 200 to 8,000 Hz. Using this initial dataset, we trained the baseline NN.

Subsequently, we generated five sets of random parameters values for the FE simulator and obtained the corresponding simulation results from the FE stimulator model (one set is shown as example in Fig. 2B, dark green dots). These spectra were then fed into the trained NN, which returned probability distributions for each parameter (magenta lines in the diagonal panels in Fig. 2A). The NN also returned the maximum *a posteriori* (MAP) values for each parameter (red vertical lines in diagonal panels and red dots in off-diagonal color maps in Fig. 2A). The actual parameter values used in the FE simulator are shown as dark green vertical lines in the diagonal panels and black dots in the off-diagonal color maps.

Key observations from the SBI outcome include:

1. All MAP lines are predicted close to the actual values, all within the predicted distributions.
2. Some parameters exhibit very narrow distributions (e.g., Young’s modulus of the eardrum and joints), indicating that those parameters are highly sensitive.
3. Some parameters have broader distributions, sometimes with no clear peaks (e.g., Young’s moduli of tendons), indicating that they are the less sensitive parameters.

The color maps (Fig. 2A, small color sub plots) illustrate the pairwise marginals of the posterior distribution, demonstrating the interactions among parameters in a multi-dimensional space. Notably, the MAP values (magenta dots in heatmaps) are not necessarily at the peaks of the marginal probability distributions, as SBI considers higher-order correlations to determine the MAP values. All MAP parameter values (red dots) are predicted close to the actual values (dark green dots).

We then fed the MAP values back into the FE simulator and compared the resultant output spectra with the ground truth spectra used to train the NN (Fig. 2B, red and dark-green lines, respectively). In all cases, the predictions closely matched the ground truth.

### B. Inferring the ME parameters of the experimental dataset

After validating the trained SBI, we imported the mean experimental spectral data to the baseline trained NN to infer model parameters capable of reproducing similar responses (Fig. 3). SBI yielded probability distributions (magenta curves) and MAP estimates (vertical red lines and dots), as illustrated in Fig. 3A. The diagonal depicts 1D marginal distributions, while the upper diagonal shows 2D paired distributions. It is important to note that the diversity of the experimental data, reflected by the standard deviations across all experiments, was incorporated into the NN training as plausible noise, explained in Methods D.

The results shown in Fig. 3A indicate that there was a tendency for the SAL YM (row 5) and ligament YM (row 4) to be on the lower limits of the prior ranges, while the Young’s modulus of joints (row 3) was on the upper boundary of its prior range. The Young’s modulus of the eardrum exhibits a distinct peak, centered around 3 MPa (row 6), consistent with results from previous studies (see Introduction).

To show robustness of the estimated values, we sampled five sets of parameter values from the probability distribution and performed FE simulations using these values (Fig. 3 A, dark-green vertical lines in 1D and triangular points in paired histograms). The resulting simulated spectral data of both MAP and the 5 samples (Fig. 3B, red and dark-green curves, respectively) are consistent with each other and compare well with the objective experimental data (black lines with gray STDs).

The impact of training the SBI with each dataset independently — stapes velocity (Vst/Pec), impedance (Zec), and absorbance magnitude (Abs) vs all of three together is reported in SI-2. New NNs were trained for each data subset. This attempt was made to demonstrate how the NN performs in predicting parameter values when only a subset of experimental data might be available. For example, in previous studies (e.g., Motallebzadeh et al., 2017b), only the impedance of the ear was used to estimate FE parameters. Not surprisingly, SI-2 shows that if an FE model is tuned to reproduce, for example, impedance only, it may not be effective in predicting stapes velocity. Furthermore, SI-2D demonstrates the ability in SBI to incorporate all available data to predict parameter values of an FE model, allowing it to reproduce multiple reference datasets simultaneously.

### C. Data noise and robustness of SBI

To account for observation noise and data variability, Gaussian noise was incorporated into the simulations used to train the NN. To assess the robustness of SBI across different noise scales, we conducted experiments with noise scales ranging from 10 times smaller to 10 times larger than the experimentally measured standard deviation. The variance of this noise was derived from experimental sets of velocity, impedance, and absorbance data as 1xSTD (standard deviations) from experimental data (gray area in Fig. 3B). NNs were trained using various noise scales added to the simulator output (after Fig. 1C).

Experimental data were introduced to the trained NN that generated individual and paired histograms of parameter distributions (Fig. 4B), and MAP parameter values were subsequently imported to the FE simulator to produce spectra compared with objective data (Fig. 4A).

As anticipated, increasing the input data noise (uncertainties) by 10xSTD expanded the range of parameter distributions (Fig. 4B). The lower input data noise of 0.1xSTD slightly narrows the probability distribution of the parameters (Fig. 4A) in comparison with those in 1xSTD (Fig. 3A). Nonetheless, the predicted MAP parameter values effectively reproduced the objective data (Fig. 4C), demonstrating robustness of the NN under varying noise conditions.

### D. Effect of the number of training dataset

As detailed in the Methods section, we employed 10,000 simulations to train the baseline NN within the SBI framework. To assess the impact of training dataset size, we conducted an evaluation of the effect of decreasing the number of paired simulations (Fig. 5). NNs were trained using 100, or 1,000 datasets. This process generated individual and paired histograms of parameter distributions (Fig. 5A, B), and MAP parameter values were subsequently imported to the FE simulator to generate spectra compared with experimental data (Fig. 5C).

As expected, decreasing the size of the training dataset to 100 or 1000 sets widened the range of parameter distributions (Fig. 5A, B). The FE model response produced by MAP values showed significant improvement in matching the objective data as the dataset size increased from 100 to 1,000 (Fig. 5C, green and blue lines). However, there was little additional improvement in the FE model response in increasing the training data from 1,000 to 10,000 (Fig. 5C, blue and red lines). Nonetheless the narrower range in the histograms (Fig. 5B vs Fig. 3A) indicates that the certainty of parameter distributions improved, which indicates greater confidence in the derived parameters with larger training dataset.

## Discussion

In this study, we applied SBI, a machine-learning method, to tune a finite-element model of the human ME using multiple sets of experimental measurements simultaneously. SBI represents a departure from conventional sensitivity analysis, which is often time-consuming, labor-intensive, and subjective in nature. A key technical difference is that in this approach, multiple parameters were varied simultaneously, whereas in the conventional approach, only one parameter is typically varied at a time. Another advantage is that, unlike in traditional methods, SBI provides probability distributions of parameter values and their interactions, leading to quantitative insights into the sensitivity of measured data to model parameters. We further explored how input data noise increases the variance of the parameter distribution but their mean values and thus the FE model simulation results were not significantly affected.

### A. Efficiency of the SBI

In this study, SBI replaced the laborious and subjective tuning procedures typical of simulators with multiple input parameters, which traditionally rely on one-at-a-time sensitivity analysis. A common limitation of biological tissue simulators such as FE models, is their reliance on single parameter sets, despite achieving similar results with different parameter combinations. SBI addresses this by returning probability distributions of parameters rather than single for values simultaneously satisfying multiple objectives, thereby narrowing down ranges capable of replicating objective data. It explores parameter combinations in multiple dimensions to derive maximum *a posteriori* (MAP) values. Ebrahimian (2023) explored sensitivity in deterministic models to input parameter variations, yet lacked inherent system randomness. SBI, while deterministic in model structure, employs probabilistic parameter distributions, enhancing model robustness as a stochastic framework.

Another significant advantage of SBI is its ability to incorporate noise and uncertainties present in experimental reference data during training. Uncertainty levels, quantified by randomized standard deviations (STDs) in our study, were integrated into simulation results of a deterministic model. We demonstrated that our trained NN effectively handled input noise variability up to 10 times the STD of the experimental data, while maintaining robust parameter probability distributions.

Global search methods typically aim to identify a single best-fitting parameter set based on measures of similarity, such as mean-squared error, between experimental and simulated data. This is different from the neural network trained by SBI that minimizes the Kullback-Leibler divergence (KL-D) between the true posterior and the approximate NN posterior for any observed data (Papamakarios and Murray, 2016). Therefore, the loss of SBI is minimized when the returned probability distribution precisely matches the Bayesian posterior probability distribution.

### B. Model parameters and geometrical effects

As described in the Methods section, the model’s geometry was reconstructed from a micro-computed tomography (μCT) imaging scan of a temporal bone used for the physiological measurements. Soft tissues and their boundaries are often not well defined from μCT scans, necessitating manual segmentation of fine soft-tissue structures. The dimensions of small ME components, such as ligaments and eardrum thickness, may be comparable to voxel dimensions (e.g., 15.1 μm), leading to potential over- or under-estimation in segmented portions of the images.

Stiffness is proportional to both the thickness and Young’s modulus. In this study, SBI predicted the distribution of Young’s modulus for the stapedial annular ligament (SAL), on the order of kilo-Pascals, which was on the lower limit of the prior range (see Fig. 3) —a rarity for fibrous tissues (Gan et al., 2011). This indicates that the SAL thickness may have been overestimated during segmentation, which was compensated by a lower Young’s modulus.

Conversely, the Young’s modulus of the joints exhibited the opposite trend. It appears that their geometrical dimensions were underestimated in segmentation, leading SBI to infer a higher Young’s modulus compared to the eardrum, despite fibrous tissues like the eardrum typically has a higher Young’s modulus.

### C. General considerations

In addition to geometric considerations discussed in the previous section, applying SBI to tune models against experimental data involves balancing number of parameters and accuracy of the simulator. This study was simplified by neglecting the inherent viscoelastic behavior of tissue to reduce model parameters. For instance, the eardrum’s effective Young’s modulus varies significantly with frequency (Daphalapurkar et al., 2009), potentially impacting its mechanical response. Another simplification involved representing the cochlear load with a viscous force on the stapes footplate to reduce the computational expense required for a full cochlear model (Kim et al., 2014).

All material parameters assumed isotropy, despite evidence of anisotropy in structures like the eardrum (Fay et al., 2006), influencing stiffness mechanisms across frequencies. The 7 parameters that we varied are also associated with ME pathologies. For example, ossicular fixation pathology is associated with high Young’s modulus of ligaments calcification and stiffening. Parameters like the stiffness of incus ligaments and tendons were consolidated due to minimal expected influence on stapes velocity or ME absorbance and impedance (Fig. 2). Understanding the influence of parameters on ME impedance and transmission often relies on prior knowledge or systematic sensitivity analysis within parameter ranges.

The variability in experimental data stems from multiple factors including background noise, tissue conditions (e.g., tissue dehydration), and positioning of measurement instruments (Chien et al., 2006). Despite efforts to maintain consistency, factors like microphone placement and laser angle can vary experimentally, affecting data reproducibility. An example of how the data extraction location in the FE model affects results is shown in SI-1. If pressure is calculated at an inaccurate location, the simulation data may never match the reference experimental data, regardless of parameter values, preventing the SBI from converging to accurate predictions. Therefore, a key criterion for successful training of the SBI NN is ensuring that simulation data from random parameter values covers the experimental data across the relevant frequency range.

The SBI toolkit allows for parameter inference through iterative loops, adjusting parameter values fed into the simulator to refine model-specific training with fewer initial training sets. However, in this study, we used pre-simulated data: a large dataset of 10,000 parameter values, each randomly sampled within predefined prior ranges, fed into the FE model (simulator). These paired inputs and outputs were then used to train the NN in a single batch. This method offers several advantages. Once simulations and training are completed, predicting parameter values for new experimental data becomes rapid and does not necessitate additional simulations or training. Moreover, utilizing pre-simulated data allowed us to investigate internal NN parameters such as the density-estimator algorithm and the impact of noise levels. It is noteworthy that, while generating the 10,000 simulation results took several weeks, training the neural network required less than 30 minutes.

## IV. Conclusions

In this study, we incorporated a novel application of SBI to tune a finite-element model of the human middle ear. SBI learns the association between model parameters and outcomes, and when presented with reference data, it provides probability distributions of the parameter values that can reproduce similar responses. This approach is practical for tuning parameters of computational models in hearing science in general, addressing both systematic and contingent uncertainties, if those parameters can be represented quantitatively. SBI is an objective method that substitutes for subjective and laborious systematic sensitivity analysis, providing not only probability distributions for each parameter (in contrast to single values) but also determining the interactions between parameters. While our findings and discussions are specific to the human middle-ear model used in this study, the method is applicable to analogous computational models, particularly those in biological contexts.

## Acknowledgements

This work was supported in part by the National Institute of Health (NIH/NIDCD R21DC020274), the Canadian Institutes of Health Research (PJT-189955), the German Research Foundation (DFG) under Germany’s Excellence Strategy (EXC 2064 – Project number 390727645) and the German Federal Ministry of Education and Research (Tübingen AI Center, FKZ: 01IS18039A). MD is a member of the International Max Planck Research School for Intelligent Systems (IMPRS-IS).

