## Supporting information for "From Simulations to Inference: Using Machine Learning to Tune Patient-Specific Finite-Element Models of the Middle Ear Towards Objective Diagnosis"

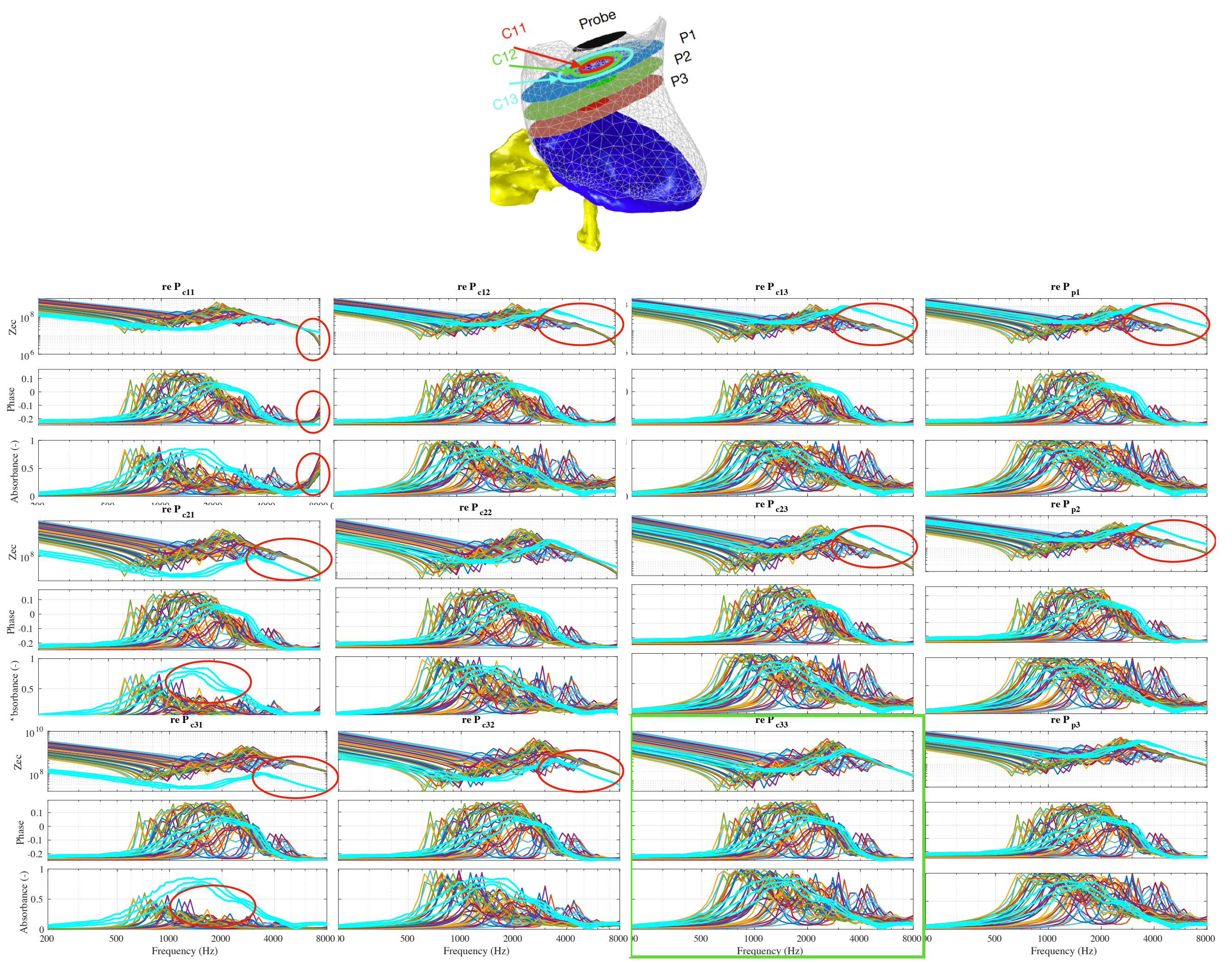


**SI-1 Effect of data extraction location on the FE model.** To calculate ear canal input impedance and absorbance, acoustic pressure and volume velocity were measured in three cross-sections at planes 1 mm, 2 mm, and 3 mm (P1-P3, respectively) parallel to the probe tip surface (indicated by the black circle in A). At each plane, measurements were taken on three concentric circles with radii of 1 mm, 2 mm, and 3 mm, resulting in a total of nine measurement locations (CXY, where X is the plane number and Y is the radius of the circle). The cyan lines represent the experimental data (mean ± standard deviation), and the colored lines represent 100 samples of the training data. Only in C33 do the simulation data cover with the experimental data; in the other locations, systematic deviations occur (depicted by red circles) regardless of material parameters.

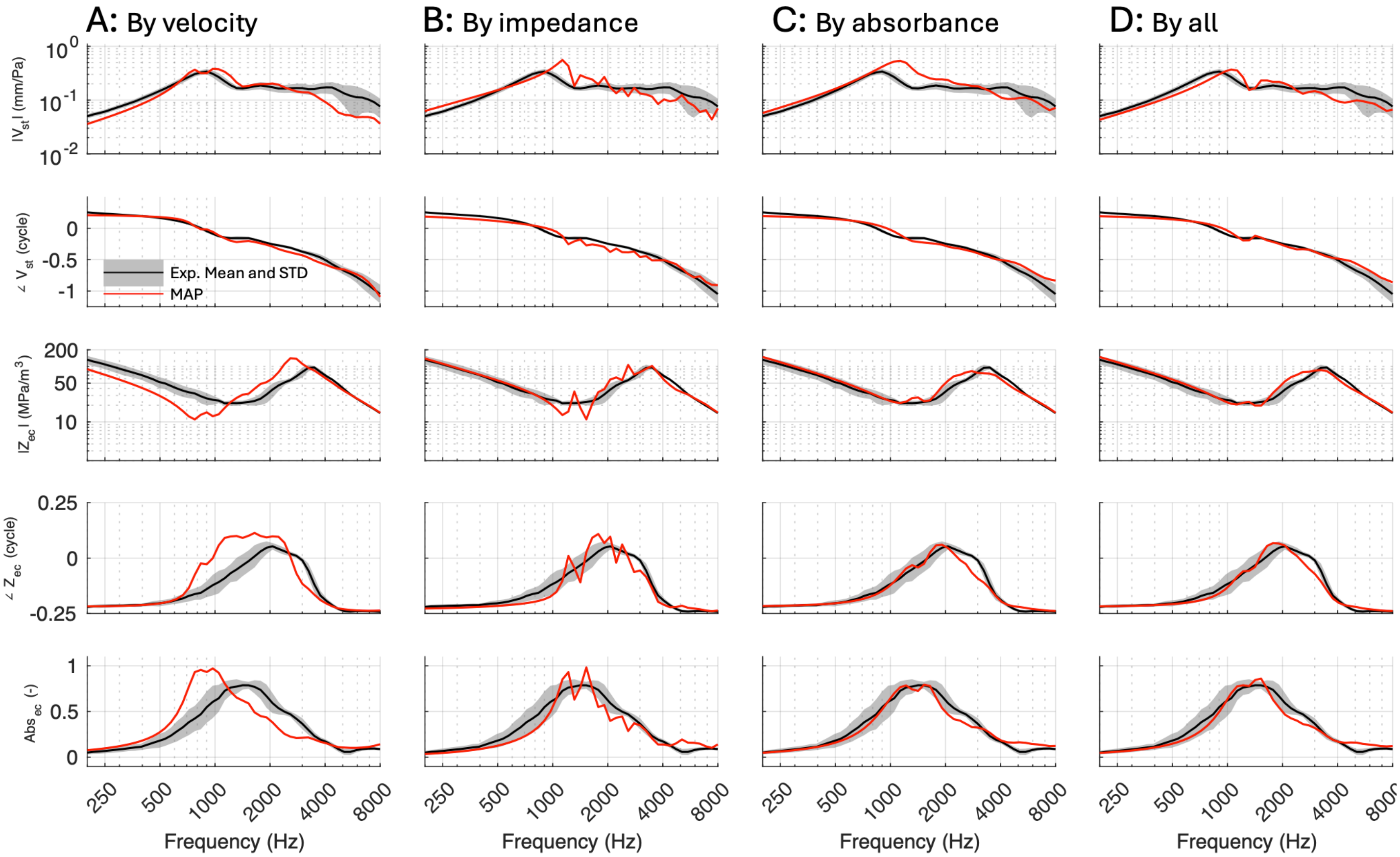


**SI-2: Training Neural Networks with Individual or Combined Datasets.** Three neural networks (NN) were trained using simulation data from each dataset individually (A: trained only on velocity, B: trained only on impedance, C: trained only on absorbance, D: trained on all datasets simultaneously – baseline NN from Fig. 3). The maximum a posteriori (MAP) values from each NN were imported into the finite element simulator, and the resulting spectra were compared with experimental data. Each NN could reproduce the spectra corresponding to its training dataset but failed to accurately reproduce spectra from other datasets. In D, when all three datasets were used to train the NN, the resultant MAP values accurately reproduced all three objectives.
